# Chromosome-scale reference genome of *Pectocarya recurvata*, a species with one of the smallest genome sizes in Boraginaceae

**DOI:** 10.1101/2024.10.14.617638

**Authors:** Poppy C. Northing, Jessie A. Pelosi, D. Lawrence Venable, Katrina M. Dlugosch

## Abstract

**Premise:** *Pectocarya recurvata* (Boraginaceae), a native species of the Sonoran Desert, has served as an important model system for a suite of ecological and evolutionary studies. Despite its relevance as an eco-evolutionary model, no reference genome assemblies in the Cynoglossoideae subfamily have been published.

**Methods:** Using PacBio HiFi sequencing, we assembled a reference genome for *P. recurvata* and annotated coding regions with full-length transcripts from an Iso-Seq transcriptome library. We assessed genome completeness with BUSCO and used flow cytometry and K-mer analysis to estimate the genome size of *P. recurvata*.

**Results:** The chromosome-scale reference genome assembly for *P. recurvata* was 216.0 Mbp long with a contig N50 of 12.1 Mbp. Our assembly included 12 primary contigs bounded by telomeres at all ends but one, consistent with the 12 chromosomes documented for the species. The chromosomes covered 158.3 Mbp and contained 30,655 predicted genes. Our measured haploid genome size from the same population was 386.5 Mbp, among the smallest for Boraginaceae. Genomic analyses suggested that this may reflect a recent autotetraploid, such that predicted diploid genome size would be even smaller and similar to the assembly size.

**Discussion:** The *P. recurvata* assembly and annotation provide a high-quality genomic resource in a sparsely represented area of the Angiosperm phylogeny. Our new genome will enable future ecophysiology, biogeography, and phylogenetics research.

## INTRODUCTION

*Pectocarya recurvata* I.M. Johnst is a winter annual forb in the Cynoglossidae subfamily and Amsinckiinae subtribe of Boraginaceae (Boraginales; Figure 1; Chacón et al., 2016; Luebert et al., 2016; Simpson et al., 2017a). *Pectocarya recurvata* is distributed across the Sonoran, Chihuahuan, and Mojave deserts of Northern Mexico and the American Southwest, regions known for unpredictable winter precipitation regimes (Noy-Meir, 1973; Huxman et al., 2004). As a winter annual plant in this region, *P. recurvata* relies on cues from late fall and early winter rains to appropriately time its germination, growth, and reproduction to take advantage of the cooler, wetter conditions that occur immediately after rain events (Mulroy and Rundel, 1977; Huxman et al., 2008). The Sonoran desert winter annual plant community, including *P. recurvata*, has served as an important model system in understanding how species adapt to variable environments and subsequently how these environments promote species coexistence (Pake and Venable, 1995, 1996; Chesson et al., 2004; Angert et al., 2009; Huxman et al., 2013). These investigations have been largely facilitated by the long term vegetation plots at the Desert Laboratory at Tumamoc Hill (Tucson, Arizona, USA), at which detailed vital rates for the most abundant species (including *P. recurvata*) have been recorded since 1982 (Venable, 2007; Huxman et al., 2013). Despite its importance to long term eco-evolutionary studies, there is no published genome or transcriptome for *P. recurvata* or any species in the Cynoglossidae subfamily.

**Figure 1.**
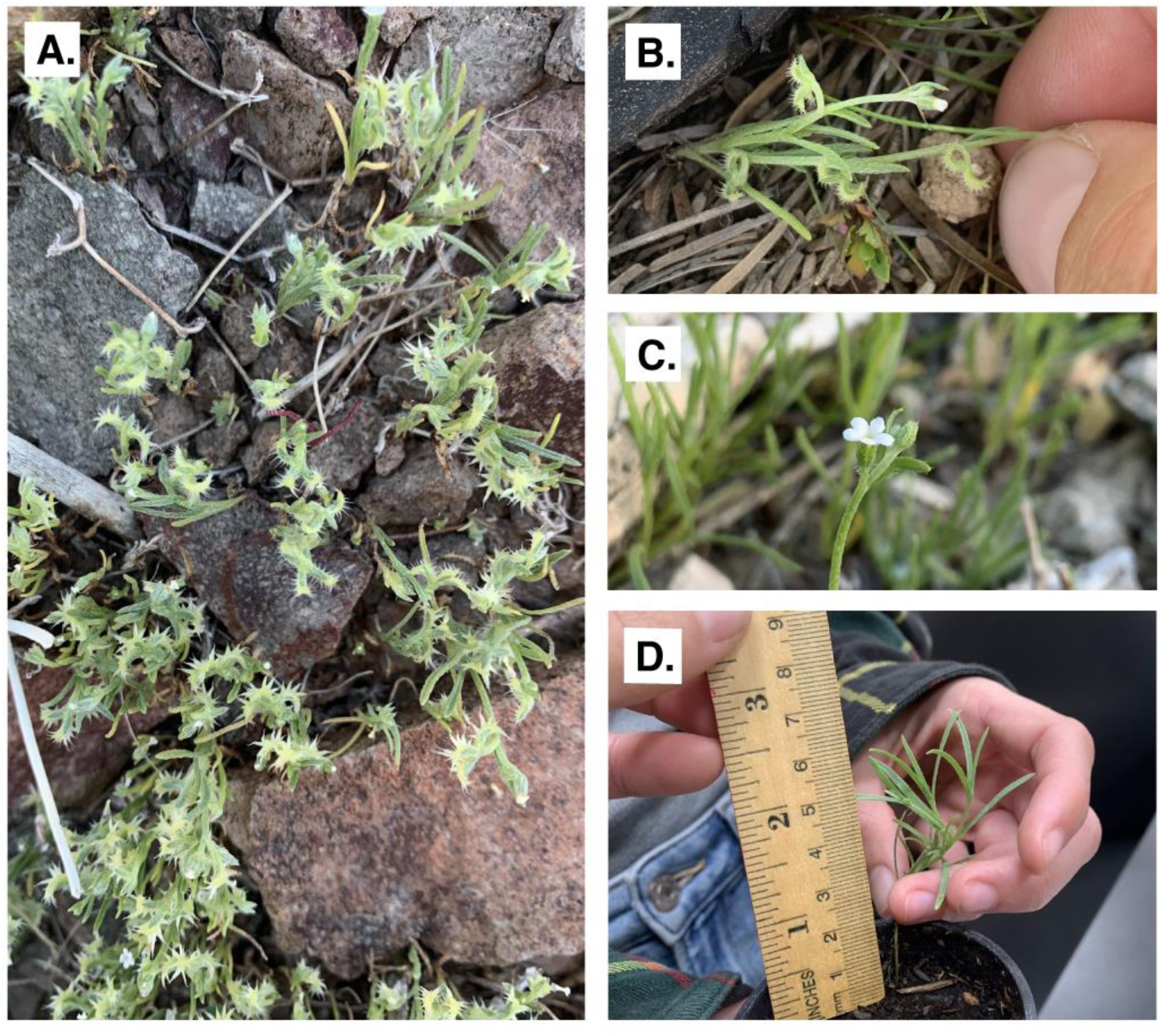
Morphology of *Pectocarya recurvata*. **A)** Individuals growing amongst individuals of *P. platycarpa* in the crevices of small rocks near Lake Havasu City, Arizona. **B)** *P. recurvata* with many fruits (four fused nutlets). **C)** A typical *P. recurvata* flower. **D)** The individual sampled for genomic and transcriptomic sequencing (ARIZ 444648). Photos taken by Poppy C. Northing.

Stable coexistence within the Sonoran desert winter annual plant community is maintained through temporal variability in precipitation and temperature acting on variation in a fundamental physiological tradeoff between growth and reproduction (Angert et al., 2007, 2009; Huxman et al., 2008). Species are separated along a specific tradeoff between water use efficiency (the amount of carbon fixed per unit of water lost) and relative growth rate (Angert et al., 2007; Huxman et al., 2008). As a more resource-conservative member of this community, *P. recurvata* exhibits a slow growth rate and high water use efficient physiology, which is associated with the maintenance of photosynthesis at lower temperatures to optimize growth after winter rainfall events (Huxman et al., 2008; Barron-Gafford et al., 2013). The genetic architecture underlying the physiological tradeoff between relative growth rate and water use efficiency and the potential for adaptive responses to climate change in these traits are an area of active interest in this system (Kimball et al., 2010; Angert et al., 2014).

More broadly, taxa in Boraginaceae have served as models to investigate the biogeographical processes underlying commonly observed disjunctions in plant distributions (Gottschling et al., 2004; Moore et al., 2006; Guilliams et al., 2017; Mabry and Simpson, 2018). Several genera in the Amsinckiinae subtribe, including *Pectocarya*, contain species that are distributed across the subtropical and temperate regions of North and South America (but not within the tropics themselves), a distribution pattern referred to as the American amphitropical disjunction (AAD; Raven, 1963; Guilliams et al., 2017; Simpson et al., 2017b). It is inferred that long distance dispersal events are the driver of AAD distributions (Raven, 1963; Wen and Ickert-Bond, 2009), yet the types of propagule morphology that might promote long distance dispersal and the features associated with successful establishment after colonization are less clear (Nathan, 2006; Chacón et al., 2017; Harris et al., 2018). In Amsinckiinae alone, at least 18 distinct long distance dispersal events across the American tropics have taken place in the relatively recent past (Chacón et al., 2017; Guilliams et al., 2017).

In Boraginaceae, there are currently (as of September 2024) three publicly available genome assemblies: *Lithospermum erythrorhizon* Siebold & Zucc. (Auber et al., 2020), *Echium plantagineum* L. (Tang et al., 2020), and *Pentaglottis sempervirens* (L.) Tausch ex L.H.Bailey, which are all taxa placed within the Boraginoideae subfamily. Cynoglossoideae is the largest subfamily of Boraginaceae, containing over 1000 species across 50 genera (Chacón et al., 2016), yet no genome assemblies are currently available from this branch of the Boraginaceae phylogeny. At the same time, the relationships of the major orders that comprise the core Lamiidae phylogeny (Boraginales, Gentianales, Solanales, Lamiales) remain unresolved (Bremer et al., 2002; Soltis et al., 2011; Zhang et al., 2020). The majority of core Lamiidae phylogenies have been constructed with plastid or mitochondrial genes (but see Zhang et al., 2020). Expanding the availability of nuclear genomic resources into additional areas of the Lamiidae phylogeny could help clarify the relationships between these orders.

Here, we present a chromosome-scale genome assembly for *Pectocarya recurvata,* the first reference genome in the Cynoglossidae subfamily of Boraginaceae. The *P. recurvata* genome assembly is a near telomere-to-telomere chromosome level assembly accompanied by structural and functional gene annotation as well as a complete chloroplast genome assembly and annotation. We also report the smallest genome size estimate to date for a species in Boraginaceae (reported C-values range from 270 – 16,000 Mbp), as well as putative ploidy variation in *P. recurvata*. Additionally, we find evidence of a whole genome duplication (WGD) in the evolutionary history of *P. recurvata* congruent with previously described WGDs in Boraginaceae (Ren et al., 2018; Tang et al., 2020). We use gene synteny analysis to compare genome structure between *Echium plantagineum* and our assembled *P. recurvata* chromosomes. Our annotated *P. recurvata* genome and plastome assemblies provide essential genomic tools for future opportunities to investigate outstanding questions in physiological ecology, biogeography, and phylogenetics.

## METHODS

### Species description

*Pectocarya recurvata* is a small herbaceous forb with basally branching, strigose stems arranged with alternate, strigose to bristle-haired linear leaves (Fig 1A; Veno, 1979; Guilliams and Kelley, 2021). It has racemose inflorescences consisting of self-pollinating flowers with small, white, funnelform corollas and seeds are fertilized autogamously (2-3mm wide; Fig 1C). *Pectocarya recurvata* is distinguished from other species in the genus by its characteristic fruit, which are four fused, recurved nutlets with toothed margins that enable epizoochoric dispersal (Fig 1A-B; Veno, 1979; Guilliams and Kelley, 2021). Within its range spanning the Chihuahuan, Mojave, and Sonoran deserts of the Southwestern United States and Northwestern Mexico, *P. recurvata* commonly occurs in patches at low elevation (10-1600 m), often growing abundantly in the shelter of rocks and larger plants such as *Larrea tridentata* (DC.) Coville (Creosote; Figure 1A; Guilliams and Kelley 2021). The haploid chromosome number of *P. recurvata* is 12, counted by Veno (1979) with meiotic chromosome squashes using individuals from five, widely distributed, populations.

### Genome size estimation

The genome size of *P. recurvata* was estimated using flow cytometry with a FACSCanto II flow cytometer (BD Biosciences, San Jose, California, USA). The 2C DNA content was measured from four individuals grown from seed sourced from the Desert Laboratory at Tumamoc Hill (32°13′ N, 111°01′ W; Tucson, Arizona, USA) in 2016. Measurements took place on three different days. The estimated haploid genome size (1C) for each sample was calculated from the 2C DNA content following the formula described by (Dolezel and Bartos, 2005).

Sample preparation and analysis by flow cytometry was conducted following a modified 2-step protocol described in (Dolezel et al., 2007) and (Cang et al., 2024). In brief, nuclei were isolated from approximately 20 mg of freshly collected leaf tissue from each *P. recurvata* sample and from *Raphanus sativus ‘saxa’* (2C DNA content = 1.11pg), which was used as an internal size standard. *Raphanus sativus* seed was provided by the Institute of Experimental Botany (Prague, Czech Republic; Dolezel et al., 2007). Leaf tissue from both species was chopped simultaneously on an ice-cold glass petri dish in 200 mL of ice cold Otto I buffer, using a new, sharp razor blade for each plant. The nuclear suspension was filtered through 18um Nylon mesh into a flow cytometry tube and kept on ice. Subsequently, 280 mL of Otto II buffer was added to the suspension, followed by 20 uL of room temperature 1 mg/ml RNase and 23 uL of ice cold 1 mg/mL propidium iodide (PI) staining solution. Each sample was then gently vortexed and incubated for 5 minutes before analyzing on the FACSCanto II with a medium flow rate.

### Reference tissue sampling

Plant tissue for the reference genome was sourced from a *P. recurvata* seed collected in the spring of 2019 at the University of Arizona’s Desert Laboratory at Tumamoc Hill (Tucson, Arizona, USA). The seed was germinated in soil in a D25L deepot (Steuwe and Sons, Tangent, Oregon, USA) at 10 ℃ and reared in a greenhouse at the University of Arizona (Tucson, Arizona, USA) in ambient light conditions and hand watered twice daily. After 77 days, the individual had produced several pairs of true leaves and 0.21 g of stem and leaf tissue was collected (Figure 1D), flash frozen in liquid nitrogen, and stored at –80 ℃ at the Arizona Genomics Institute (AGI); this tissue was used to prepare genomic sequencing libraries (see below). The remaining root, stem, and leaf tissue was returned to the same greenhouse and continued growing. After 66 days of additional growth, 0.52 g of leaf, stem, and reproductive tissue was flash frozen in liquid nitrogen and stored at –80 ℃ at the AGI; this tissue was used to prepare the transcriptome sequencing library (see below). A reference voucher of the remaining plant tissue from this individual is archived in the University of Arizona herbarium (ARIZ 444648).

### Genome and transcriptome sequencing

Genome sequencing was conducted by the AGI following PacBio’s HiFi protocols for library preparation and sequencing. After high molecular weight DNA was extracted using the protocol described by Doyle and Doyle (1978) with minor modifications and assessed for quality control with a Femto Pulse system (Agilent, Santa Clara, California, USA), DNA was sheared to 10-20 kb with a Megaruptor 3 sonicator (Diagenode, Denville, New Jersey, USA) and treated with SMRTbell cleanup beads (PacBio, Menlo Park, California, USA). The final HiFi libraries were constructed using a SMRTbell Prep Kit v3.0 (PacBio), size selected to 10-25 kbp using a PippinHT size selector (Sage Science, Beverly, Massachusetts, USA), and validated with the Femto Pulse system (Agilent). This final, size-selected library was prepared for sequencing using the PacBio SequelII Sequencing Kit v3.1 for HiFi libraries, loaded onto one PacBio Revio SMRT cell, and sequenced for 24 hours in circular consensus sequencing (CCS) mode (PacBio), yielding 65.6 Gbp of HiFi sequencing data.

An Iso-Seq library was also constructed by the AGI to enable gene identification and annotation of the *P. recurvata* genome assembly. Total RNA was isolated from the flash frozen sample of pooled leaf, stem, and reproductive tissue with PureLink® Plant RNA Reagent kit (Thermo Fisher Scientific Inc., Waltham, Massachusetts, USA) and the RNA Integrity Number (RIN) was measured for quality control using an Agilent 2100 Bioanalyzer (Agilent; the sample RIN = 7.8). The Kinnex Iso-Seq library was prepared by first synthesizing cDNA from 300 ng of total RNA using Iso-Seq express 2.0 kit (PacBio) and the size distribution of the resulting cDNA was checked with a Bioanalyzer (Agilent). The final Iso-Seq library was constructed following the PacBio Kinnex PCR protocol (PacBio) and sequenced on one Revio SMRT cell, generating 75.7 Gbp of HiFi reads comprising 85 million transcripts that reduced to 1.65 million isoform transcripts after clustering analysis was done using the *IsoSeq* software toolkit (https://isoseq.how/, PacBio).

### Assembly

A kmer analysis of the HiFi reads was used to estimate genome size, abundance of repetitive elements, heterozygosity, and ploidy using KMC v3.1.0 (Kokot et al., 2017), GenomeScope v1.0 (Vurture et al., 2017), and SmudgePlot v1.0 (Ranallo-Benavidez et al., 2020) with a maximum kmer coverage of 10,000 to count and analyze 17-mers, respectively. The initial genome assembly was constructed de novo from the PacBio HiFi reads using Hifiasm v0.16.1 (Cheng et al., 2021). Based on low estimated heterozygosity from the kmer analysis (see Results) and the selfing mating system of *P. recurvata*, purging of haplotypic duplications was skipped in the assembly (using the *-l0* flag). The resulting assembly graph was visualized using Bandage v0.8.1 (Wick et al., 2015) and subsequently evaluated and error-corrected using Inspector with the –-pacbio-hifi flag (Chen et al., 2021). The initial assembly was assessed for completeness using 2326 Benchmarking Universal Single-Copy Orthologs (BUSCOs) from the Eudicots dataset *eudicots_odb10* (01 Aug 2024) using BUSCO version 5.6.1 (Manni et al., 2021).

The initial assembly consisted of 12 large contigs and 6,449 smaller, trailing contigs (see *Results*; **Table 1**). Three approaches were taken to identify the sequence content of these contigs. First, the trailing contigs were mapped to our assembled *P. recurvata* reference plastome (see below) using Geneious prime 2024.0.5 (https://www.geneious.com), and any contigs aligning to the reference plastome were removed from the assembly. Second, mitochondrial DNA sequences from species in Boraginaceae publicly available in the NCBI nucleotide database (accessed June 2024) were mapped to the trailing contigs to identify contigs containing mitochondrial DNA. Finally, the trailing contigs were mapped to the 12 large contigs to identify any repetitive genomic sequences in the contigs. While the contigs that mapped to the chloroplast were removed from the initial assembly (generating the final assembly), the contigs mapping to mitochondrial sequences and those which mapped to the genome were retained in the final assembly due to ambiguity in their placements.

**Table 1.**
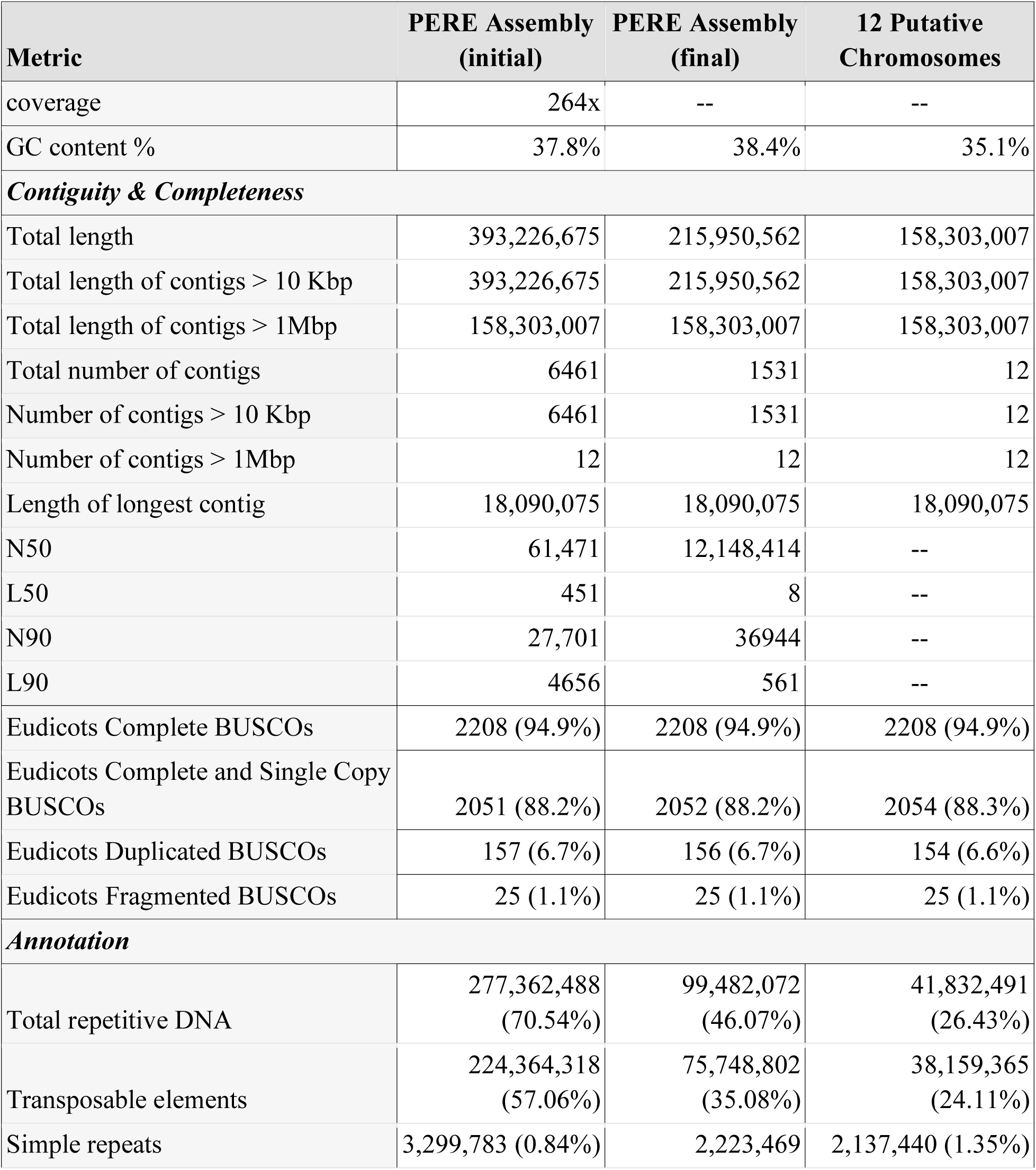
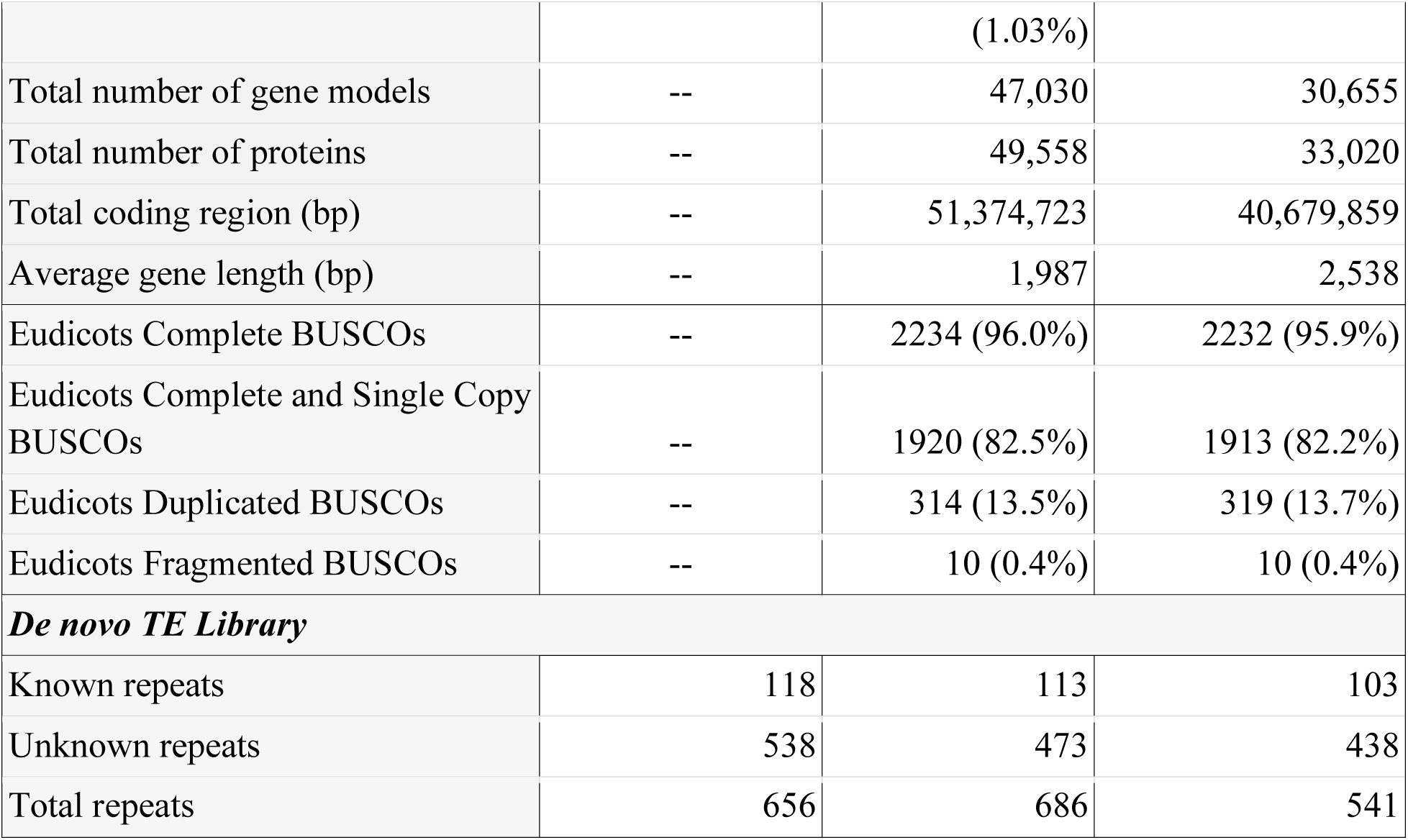
*Pectocarya recurvata* assembly and annotation statistics.

The chloroplast genome was constructed de novo using the PacBio HiFi reads. Potential chloroplast reads were identified by aligning the raw HiFi reads to the *Echium plantagineum* (Boraginaceae) reference plastome (Genbank accession: OL335188.1; Carvalho Leonardo et al., 2022) using Minimap2 v2.28 with default settings (Li, 2018). Reads that did not align to the reference were filtered and removed using Samtools v1.10 (Li et al., 2009). The remaining mapped reads had an exceptionally high estimated coverage (159,683x, see Results); accordingly, reads were randomly subsampled with SeqKit v2.8.1 (Shen et al., 2016) to approximately 150x coverage to mitigate assembly errors. These subsampled reads were then used to generate a circular *P. recurvata* chloroplast genome assembly using Hifiasm v0.16.1 with default settings (Cheng et al., 2021). The assembly was visualized and evaluated using Bandage v0.8.1 (Wick et al., 2015) and Geneious Prime 2024.0.5 (https://www.geneious.com).

### Annotation

Repetitive elements were identified, classified, and soft masked in the final assembly. First, a custom species-specific library of identified and classified repetitive element families was generated with RepeatModeler v2.0.3 using default parameters (Flynn et al., 2020), which identifies known and unknown repetitive elements de novo using RECON v1.08 (Bao and Eddy, 2002) and RepeatScout v1.0.6 (Price et al., 2005). Distinct repeats were identified and soft masked (using the *xsmall* flag) in the assembly iteratively with RepeatMasker v4.1.3 (https://www.repeatmasker.org/RepeatMasker/), starting with simple repeats (specifying the *noint* flag), then repeats identified from the *nrTEplants2020* curated repetitive sequence library (Contreras-Moreira et al., 2021), followed by known and unknown repeats from the *P. recurvata* repeat library. After repeats were masked in the final assembly, telomeric regions of the chromosomes were predicted using the TeloExplorer program from QuarTeT v1.1.6 with *-m* 85 (Lin et al., 2023).

Gene structural annotation of the *P. recurvata* genome assembly was performed on the soft-masked assembly using the BRAKER3 annotation pipeline (Hoff et al., 2016, 2019; Brůna et al., 2021; Gabriel et al., 2021, 2024), which uses evidence from homologous proteins and transcripts to train AUGUSTUS (Stanke et al., 2006) and GeneMark-EP+ (Lomsadze et al., 2005; Brůna et al., 2020) to produce a final set of high-confidence gene models. First, whole transcripts from the Iso-Seq library were aligned to the *P. recurvata* reference assembly using Minimap2 v2.28 with *-ax splice:hq* specified (Li, 2018). A custom protein database containing 456,224 peptide sequences was constructed by combining the proteomes of ten species in the Lamiids clade with well-annotated genomes (Appendix S1; see Supporting Information with this article). We used the *braker3_lr* singularity container, which accommodates the use of the *P. recurvata* Iso-Seq transcriptome as evidence (along with the custom protein database). The resulting set of predicted proteins was assessed for completeness against the *eudicots_odb10* (08 jan 2024) benchmarking ortholog database with BUSCO v5.6.1 in *euk_tran* mode (specifying *-m* ‘tran’; Manni et al., 2021). Summary statistics describing the protein annotation were generated using the agat_sp_statistics.pl script from AGAT v1.0.0 (Dainat, 2022). The final protein set was functionally annotated using InterProScan v5.66-98.0 with the –-goterms flag to include GO terms in the annotation (Jones et al., 2014).

The chloroplast assembly was annotated using GeSeq v2.03 (Tillich et al., 2017). GeSeq utilizes HMMER v3.4 (hmmer.org), Chloë (Zhong, 2020), and Blast (Altschul et al., 1990) to annotate the inverted repeat (IR) regions, protein coding sequences, rRNAs, and tRNAs in plastid genomes. The figure depicting the annotated *P. recurvata* chloroplast genome was generated using OGDRAW v1.3.1 (Greiner et al., 2019).

### Synteny analysis

Regions of conserved gene synteny were identified among the assembled *P. recurvata* chromosomes and with those of *Echium plantagineum* (Boraginaceae; GCA_003412495.2), the only other chromosome-level assembly in Boraginaceae that has been annotated (Tang et al., 2020). While the *Echium plantagineum* annotation described by Tang et al. 2020 is not publicly available, the raw transcripts used to annotate the reference genome are available on NCBI. Three of these available transcriptome libraries (SRR4034891, SRR4034890, and SRR7076848) were used to generate a draft gene structural annotation of the *E. plantagineum* assembly. Using the standard *braker3* singularity container, the same annotation protocol described above was followed to incorporate short-read RNA-Seq transcriptome data as evidence of gene structure (Gabriel et al., 2024). Groups of orthologous genes from the resulting proteomes from each genome annotation were identified using OrthoFinder v2.5.4 with default settings (Emms and Kelly, 2019). The groups of shared orthologs identified by OrthoFinder were used to identify blocks of conserved gene order that cluster into regions of synteny using the GENESPACE v1.4 R package with default settings (Lovell et al., 2022). The syntenic depth ratio of each syntenic block (collinear sets of genes > 5) was calculated using the pythonic version of MCscan, which depends on LAST (Tang et al., 2008; Kiełbasa et al., 2011).

Additionally, evidence for ancient whole genome duplications in the *P. recurvata* and *E. plantagineum* genomes was identified by analyzing the frequency distribution of synonymous divergence (K_s_) between paralogs (for each genome individually). Synonymous divergence was calculated for each pair of duplicate genes using wgd v2 (Chen et al., 2024). The K_s_ plot was filtered by only retaining syntenic duplicates, those duplicates that most represent duplication from a genome-wide multiplication rather than small-scale multiplications such as tandem duplicates, within wgd using i-ADHoRe v3.0 (Proost et al., 2012).

## RESULTS

### Genome assembly

The initial, Inspector-corrected Hifiasm assembly contained 6461 contigs covering 393,226,675 bp. Twelve of these contigs were greater than 1 Mbp (total length = 158,303,007 bp), which corresponds with the haploid chromosome number (n=12) of *P. recurvata* (Veno, 1979). Of the 6449 *trailing* contigs in the initial assembly, 4930 (76.4%) aligned to our *P. recurvata* plastome assembly (with 94.1% identical sequence) and were subsequently removed from the initial assembly. The Boraginaceae mitochondrial sequences from NCBI mapped to 678 (10.5%) unique contigs, and 268 (4.4%) other contigs mapped to the 12 large contigs (specifically, all mapped to the end of chromosome 12).

The final assembly contains 1531 contigs covering 215,950,562 bp and has an N50 of 12,148,414 bp and an L50 of 8, indicating a high level of contiguity (Table 1). The total length of the 12 leading contigs is 158,303,007 bp, which is similar to the estimated genome size (152.5 Mbp) from the kmer analysis and accounts for 73.3% of the final assembly. A total of 94.9% of complete eudicot BUSCOs (respectively) were found in these 12 contigs and none of the BUSCOs that were duplicated, fragmented, or missing from either BUSCO dataset were identified in the trailing contigs of the assembly (Table 1). Hereafter, we refer to these 12 leading contigs as the putative *P. recurvata* chromosomes, numbered #1-12 by length (Appendix S1).

Telomeric regions consisting of greater than 85 tandem repeats of CCCTAAA, the typical plant telomeric repeat sequence (Peska and Garcia, 2020), were identified in both ends of the putative chromosomes 1-11, while 64 repeats were identified on one end of chromosome 12 and none were found on the other end. No other telomeric regions were identified in the assembly.

The chromosomes (158.3 Mbp) consist of 26.43% repetitive content identified by RepeatMasker (https://www.repeatmasker.org/RepeatMasker/)—24.11% is attributed to transposable elements, 1.66% to tandem repeats, and 0.17% to small RNAs. The most common transposable elements are LTRs, which make up 6.29% of the chromosomes, while LINEs make up 0.81%, SINEs make up 0.22%, and 15.38% are unclassified. DNA transposons comprise 1.40% of the chromosomes (Appendix S1). Notably, the repetitive content of the *P. recurvata* chromosomes (26.43%) is much lower than that described in either of the described genome assemblies from Boraginaceae: the *L. erythrorhizon* assembly is 366.7 Mbp of which 51.78% is repetitive content, and similarly the *E. plantagineum* assembly is 348.9 Mbp and 43.30% repetitive content (Appendix S1) (Auber et al., 2020; Tang et al., 2020).

### Genome size estimation: flow cytometry, kmer analysis, and inference of ploidy

Our haploid (1C) genome size estimate of *P. recurvata* from flow cytometry was 386.5 ± 1.49 Mbp (0.395 ± 0.0015 pg). In contrast, analysis of the coverage and frequency of 17-mers from the raw HiFi reads predicted a haploid genome length of 152.5 Mbp, and our assembly of the expected 12 chromosomes had a total length of 158.3 Mbp, both just under half the size of the flow cytometry estimate. (Appendix S2; Appendix S1). The SmudgePlot generated to detect the ploidy from relative kmer coverages indicated that the individual sequenced, which was sourced from the same plots as those used for genome size estimation, was likely tetraploid (Ranallo-Benavidez et al., 2020; Appendix S2). The kmer analysis also estimated a very low level of heterozygosity (min 0.044%; max 0.051%), consistent with high levels of self fertilization and raising the possibility that our assembly represents a haploid assembly of *P. recurvata* that collapsed two, putatively highly similar, subgenomes of a very recent autotetraploid, as well as allelic variation.

### Genome annotation

The structural gene annotation of the *P. recurvata* reference chromosomes contains 30,655 predicted gene models (mean gene length = 2538 bp) that code for 33,020 proteins. On average, the predicted genes are 2538 bp long and contain 5 exons. In total, the predicted genes encompass 77,831,286 bp, of which 40,679,859 bp is attributed to coding sequence. The comparison of the *P. recurvata* protein annotation against the Eudicot odb10 BUSCO dataset found 95.5% complete BUSCOs (n=2326) and 1913 (85.7%) are single copy (Table 1). Of the 33,020 predicted proteins, 93.8% (30,979) were functionally annotated with InterProScan.

### Gene synteny analysis

Self-self comparison of gene order and content in the *P. recurvata* chromosomes revealed several duplicated collinear gene segments. The frequency distribution of synonymous substitution rate (K_s_) between paralogs in the *P. recurvata* genome revealed a distinctive peak in the number of syntenic paralogs at at K_s_ of 0.4-0.5, reflective of an ancient whole genome duplication in the evolutionary history of *P. recurvata* (Figure 2). There is also a distinctive peak at K_s_ = 0.4-0.5 in the K_s_ distribution for *E. plantagineum,* as well as a distinctive peak in syntenic paralogs at K_s_ = 0.1, reflective of an additional whole genome duplication event in the evolutionary history of *E. plantagineum* (Appendix S2).

**Figure 2.**
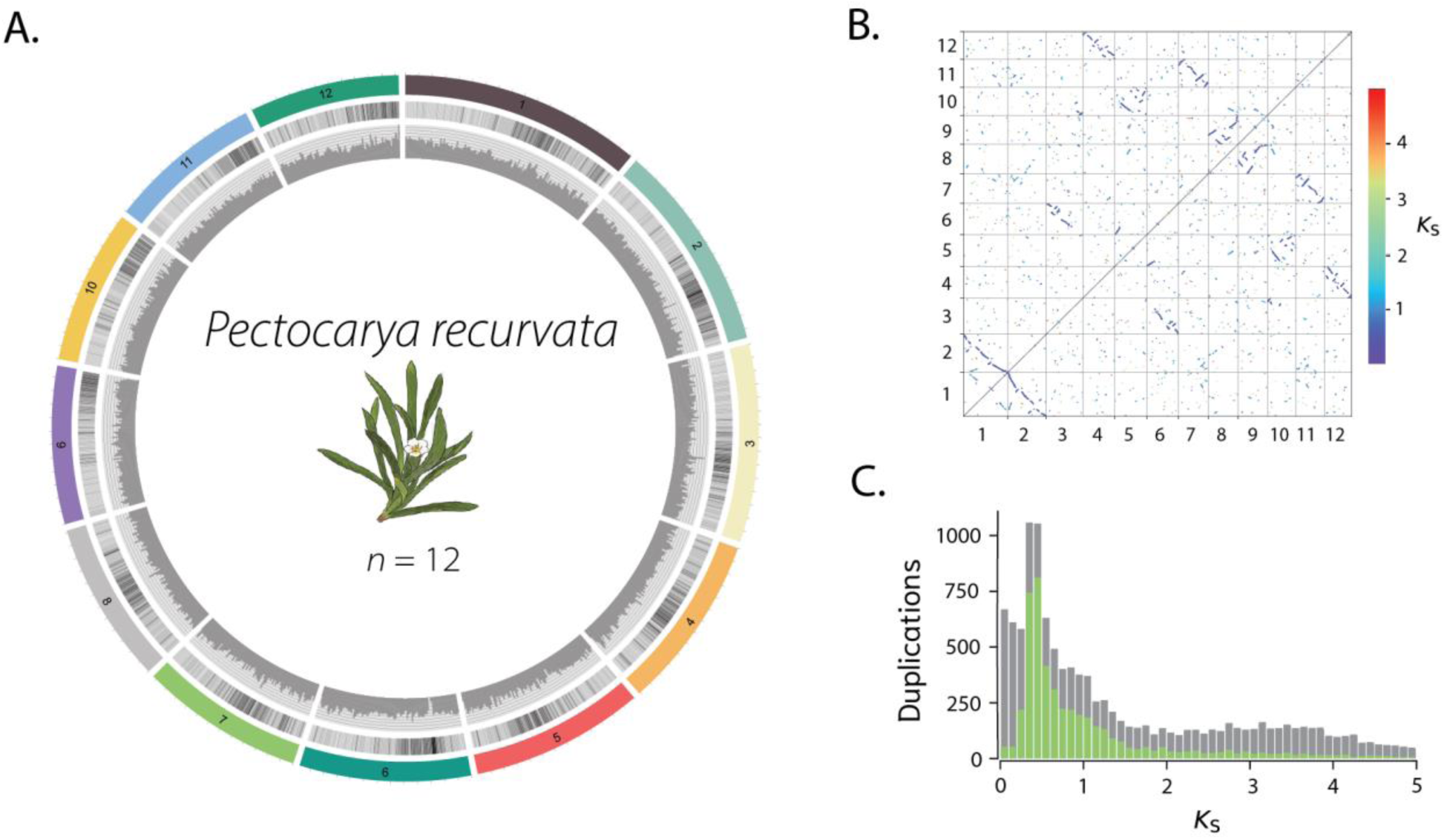
Analysis of self-self synteny with the *Pectocarya recurvata* reference chromosomes suggests a recent whole genome duplication in Boraginaceae. **A)** The assembled *P. recurvata* chromosomes (*n* = 12) labeled 1-12 by length. Transposable element density is illustrated by the grey heatmaps in the first inner circle. Gene density is depicted by the histograms in the innermost circle. Figure produced with Circos. **B)** A dotplot of self-self synteny in the *P. recurvata* chromosomes, colored according to the synonymous divergence (K_s_) between each pair of duplicated genes. **C)** A frequency distribution of the K_s_ between paralogs in the *P. recurvata* chromosome. The grey background distribution includes every paralog, while the green distribution only includes syntenic paralogs.

By comparing the gene order of the *P. recurvata* and *E. plantagineum* genomes, we identified a 2:3 syntenic relationship between the genomes of these species (Figure 3; Appendix S2). Together, the *P. recurvata* and *E. plantagineum* reference chromosomes have 70,183 annotated genes. Nearly all genes in *P. recurvata* and *E. plantagineum* were placed in 19,150 orthogroups (89.0% of all genes), of which 15,906 orthogroups were shared between the two species. We identified 6339 single copy orthologs (which account for 40% of the *P. recurvata* genome). The shared orthogroups form 1252 syntenic blocks (collinear sets of genes > 5) that cluster into 690 syntenic regions (Figure 3). Nearly half (47%) of syntenic blocks in the *P. recurvata* genome are found in duplicate in the *E. plantagineum* genome. In the *E. plantagienum* genome, 33% of syntenic blocks are found in duplicate in the *P. recurvata* genome and 28% are found in triplicate. The excess number of duplicate genes shared in both genomes suggests that these species share a whole genome duplication in their evolutionary history. The excess of triplicate genes in the *E. plantagineum* genome indicates that an additional, independent whole genome duplication has occurred in the history of *E. plantagineum*.

**Figure 3.**
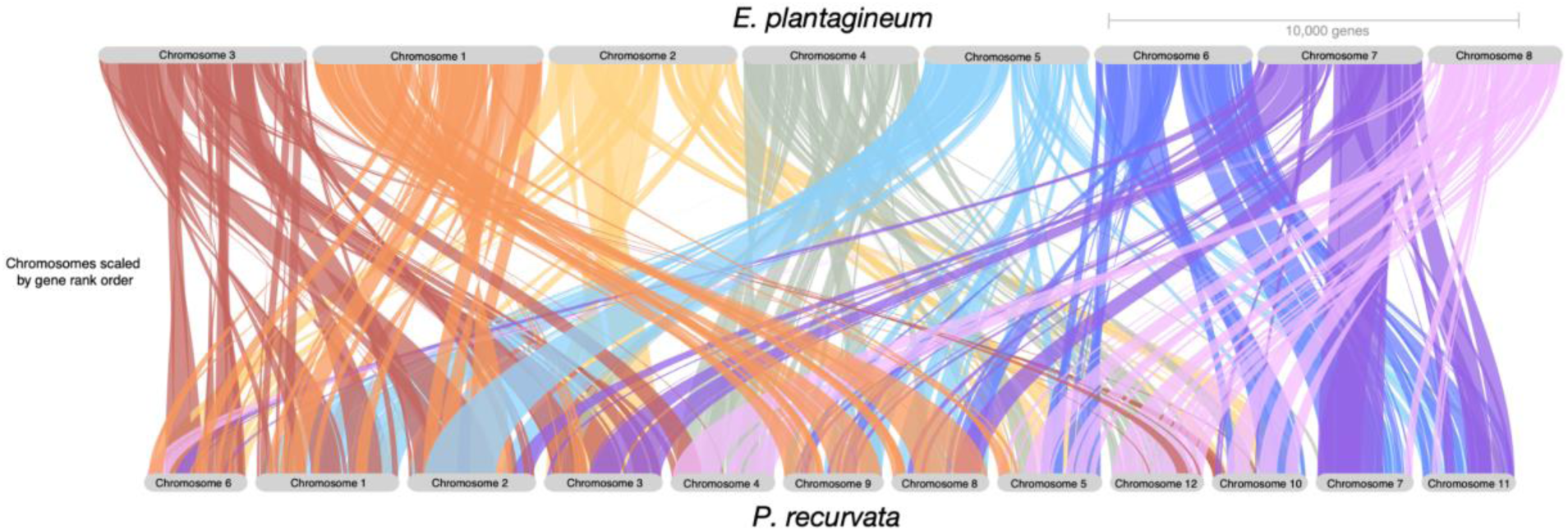
Synteny between the *Pectocarya recurvata* and *Echium plantagineum* reference chromosomes.

### Chloroplast assembly and annotation

Across 1,869,123 reads (mean length = 12,814 bp), 23,952,585,969 bp successfully mapped to the *E. plantagineum* reference plastome (149,776 bp), reflecting an estimated mean coverage of 159,683x. The initial de novo Hifiasm assembly from these reads, subsampled to 150x per-base coverage, yielded three contigs totalling 187,628 bp, consisting of one 150,172 bp circular contig and two smaller linear contigs (24,314 bp and 13,142 bp). To investigate the content of these linear contigs, both were aligned to the large circular contig using Geneious prime 2024.0.5 (https://www.geneious.com). The linear contigs each have 100% sequence identity to regions on the circular contig (the smaller contig maps to 72,063-85,205 bp; the larger 133,855-7,997 bp). As the content of these small linear contigs is 100% identical to the large circular contig, these were removed from the assembly. Thus, the final *P. recurvata* chloroplast genome (hereafter ‘plastome’) assembly consists solely of the large circular contig from the initial assembly.

The *P. recurvata* plastome has a 37.4% GC content and a quadripartite structure typical of chloroplast genomes, consisting of a small single copy region (17,143 bp; 31.1% GC), two inverted repeats (IRa: 25,446 bp; IRb: 25,467 bp; 43.1% GC), and a large single copy region (82,096 bp; 35.3% GC). GeSeq annotation of this circular contig predicted 130 unique genes in the plastome including 8 rRNAs, 37 tRNAs, and 85 protein coding genes (Appendix S2). The length and gene content of the *P. recurvata* plastid genome assembly are comparable to that of the *E. plantagineum* plastome (Appendix S1; Carvalho Leonardo et al., 2022).

## DISCUSSION

We assembled and annotated the first chromosome-level reference genome of *Pectocarya recurvata*. The final genome assembly (216.0 Mbp) is highly contiguous (N50 = 12.1 Mbp) and complete (94.9% Eudicot BUSCOs identified). This completeness is reflected in the twelve reference chromosomes, which make up 158.3 Mbp of the assembly. These chromosomes have 23 out of 24 putative telomeres and contain 30,655 predicted genes that code for 33,020 proteins that comprise highly complete structural and function annotations (95.9% Eudicot BUSCOs were identified; 93.8% were assigned functions by InterProScan). Many large syntenic regions were identified between *P. recurvata* and *E. plantagineum*, further validating the quality of our gene annotation.

The frequency distribution of the synonymous substitution rates (K_s_) of putative paralogs in the *P. recurvata* and *E. plantagineum* genomes suggests that a whole genome duplication event (WGD) occurred in recent evolutionary history at around K_s_ = 0.4-0.5 (Figure 2). Our finding is congruent with a previously described WGD in the core Boraginales at K_s_ ≈ 0.417, which putatively occurred prior to the divergence of the Boraginoideae and Cynoglossoideae subfamilies in Boraginaceae (Tang et al., 2020; Chen et al., 2023). This duplication is likely the source of the large number of genes found reciprocally in duplicate between the *P. recurvata* and *E. plantagineum* genomes. While Ren et al. (2018) and Tang et al. (2020) also found evidence of another, older WGD in the core Boraginales at K_s_ ≈ 0.939, there is no distinctive peak at K_s_ = 0.9 in our analyses. Interestingly, some recent WGD analyses using a combination of available genomes and transcriptomes do not identify the more recent WGD at K_s_ ≈ 0.417 that is evidenced by the *P. recurvata* genome and other analyses (Ren et al., 2018; Auber et al., 2020; Tang et al., 2020; Zhang et al., 2020). However, Ren et al (2018) did find evidence from the *Trigonotis peduncularis* transcriptome (Myosotideae subtribe; Cynoglossoideae tribe) of a very recent WGD at K_s_ ≈ 0.13, while Tang et al. (2020) did not. At the same time, we find evidence that *E. plantagineum* has undergone a more recent, independent whole genome duplication from *P. recurvata* (Appendix S2). Whole genome duplication has played an instrumental role in the evolution the angiosperms—on average, every angiosperm has experienced 3.5 rounds of whole genome duplication (and subsequent genome re-diploidization) in their evolutionary history (One Thousand Plant Transcriptomes Initiative, 2019; McKibben et al., 2024). It remains a prominent goal in plant biology to determine the phylogenetic placement of WGDs and their role in plant evolution (Van de Peer et al., 2017; Barker et al., 2024). Additional genomic resources in Boraginales will be helpful for confidently placing WGDs and their impact on the evolution of this clade.

In estimating the genome size and sequencing the *P. recurvata* genome, we discovered putative cytotype variation in *P. recurvata* that we infer is the result of very recent autopolyploidy. Our genome size estimate from flow cytometry was 386.5 Mbp, over twice the size of our highly complete assembly of 12 chromosomes (158.3 Mbp), using plants from the same population. SmudgePlot analysis of the kmers indicated that the sequenced individual was likely a very recent tetraploid with very low divergence among duplicated kmers. We infer that the assembly collapsed the two highly similar subgenomes, which reconciles the measured and assembled genome sizes. Prior cytological work by Veno (1979) used individuals of *P. recurvata* sourced from five populations across Arizona and Southern California to establish that the haploid chromosome number of *P. recurvata* is 12. Species in *Pectocarya* follow a polyploid series of *n* = 12 (e.g. *P. recurvata*, *P. heterocarpa*), 24 (e.g. *P. platycarpa*, *P. anisocarpa*), and 36 (*P. linearis*), suggesting that whole genome duplication has played a role in the evolutionary history of these plants (Veno, 1979). Autopolyploidy represents a form of genomic variation that has important consequences for the population genetics of a species—it is hypothesized that doubling the effective population size results in relief from inbreeding depression by maintaining genome heterozygosity (Moody et al., 1993; Ronfort et al., 1998; Soltis and Soltis, 2000; Parisod et al., 2010). Autopolyploidy is common among plants, yet autopolyploid species often are unnamed because they appear morphologically identical to their diploid progenitors (Soltis and Soltis, 2000; Soltis et al., 2007; Barker et al., 2016; Lv et al., 2024). At the same time, the short-term ecological consequences of autopolyploidy events remain elusive (Stebbins, 1940; Soltis et al., 2014; Assour et al., 2024). Quantifying the geographic extent of cytotype variation in *P. recurvata* would not only lend insight into the distribution of genetic variation in this species, but also provide the opportunity to explore associations between autopolyploidy and relevant ecological factors (Servick et al., 2015; Visger et al., 2016; Barker et al., 2024).

In addition to our discovery of putative ploidy variation in *P. recurvata*, through the sequencing and assembling of the *P. recurvata* reference genome, we uncovered one of the smallest recorded genome sizes in Boraginaceae to date. There are currently 57 primary genome size estimates for taxa in Boraginaceae recorded in the Kew Plant DNA C-values Database (Release 7.1; https://cvalues.science.kew.org/ accessed September 2024) that range from 313.0 Mbp (*Echium bonnetii*) to 10,878.0 Mbp for *Alkanna leiocarpa* (mean = 1,599.07 Mbp). Additionally, a more recent comprehensive survey of genome size estimates in Boraginaceae by Kobrlová and Hroneš (2019) recorded genome sizes that range from approximately 270 Mbp for *Myosotis sylvatica* to 16,000 Mbp for *Lycopsis arvensis*. At 386.5 Mbp, our flow cytometry measurement of a putative tetraploid is already among the smallest of these measurements, with the possibility of diploids being closer to our assembled (collapsed) size of 158.3 Mbp (12 chromosomes) – 216 Mbp (final assembly, including potential mtDNA and haplotypes), which would be smaller than all currently observations in the family. Genome size can vary greatly between related species, in part due to whole genome duplication and subsequent re-diploidization (Li et al., 2021) as well as transposable element activity (Vitte and Panaud, 2005; El Baidouri and Panaud, 2013). In Boraginaceae, taxa in the Boraginoideae subfamily are more often diploid than taxa in Cynoglossoideae, yet have larger chromosomes and genome sizes, suggesting that transposable elements might have played an important role in generating the 35-fold variation in genome sizes across clades within this group (Kobrlová and Hroneš, 2019).

The adaptive significance of plants’ genome sizes remains under investigation (Gregory, 2001; Knight and Beaulieu, 2008; Veselý et al., 2012; Bureš et al., 2024). The large genome constraint hypothesis asserts larger genome sizes are maladaptive due to the physiological consequences associated with larger cell sizes (Knight et al., 2005; Veselý et al., 2020). For example, larger stomatal guard cells open and close more slowly, resulting in lower water use efficiency (Drake et al., 2013). Such small changes in physiology may compound and decrease fitness across climates (Bureš et al., 2024). Herbaceous angiosperms with larger genomes are associated with a heightened extinction risk, regardless of climate (Soto Gomez et al., 2024). At the same time, smaller genome sizes are frequently associated with faster rates of cell division that lead to faster development and flowering times (Grime and Mowforth, 1982; Bilinski et al., 2018; Cacho et al., 2021; Cang et al., 2024). Desert annual plants, such as *P. recurvata*, experience strong selective pressure to rapidly grow and reproduce after germinating to take advantage of ephemeral moisture after precipitation events (Huxman et al., 2008; Barron-Gafford et al., 2013). Past selection has favored a highly water-use efficient phenotype in *P. recurvata* relative to other members of the Sonoran desert winter annual community (Angert et al., 2007; Huxman et al., 2008). Our finding of a small genome size in this species raises the open question of whether selection might also be favoring faster development time through genome size reduction.

While the Sonoran desert winter annual plants are a well-developed study system in ecology and evolutionary biology, only a handful of reference genomes have been assembled for these plants. Of the 51 most common Sonoran desert winter annuals found at Tumamoc Hill (Tucson, Arizona, USA; see Ge et al., 2019), only *Eschscholzia california*, *Erodium texanum*, *Sisymbrium irio* (Haudry et al., 2013), and now *P. recurvata*, have reference genomes. Much is known about how species coexistence in this community is shaped by species variation in their position along a physiological tradeoff between water use efficiency and relative growth rate (Chesson, 2000; Angert et al., 2009). Yet, little is known about the genetic architecture underlying these physiological traits in these species and how genetic differences may combine to manifest these ecological dynamics (Kimball et al., 2013; Angert et al., 2014). In the context of anthropogenic climate change, it is critical to understand the genetic basis of relevant ecological traits that are being selected on by climate change to evaluate species’ extinction risk and downstream ecological consequences (Scheffers et al., 2016; Whiting et al., 2024). The Sonoran desert winter annuals have experienced overall declines in abundance over recent anthropogenic change, and simultaneously the community is shifting to favoring higher water use efficient species (Kimball et al., 2010; Huxman et al., 2013). The addition of a *P. recurvata* genome will facilitate investigations into the genetic mechanisms driving responses to climate in these plants.

## Conclusion

*Pectocarya recurvata* is an important model system for a suite of ecological and evolutionary biology studies. Our generation of a new chromosome-level reference genome assembly and annotation for *P. recurvata* opens up the possibility of new lines of inquiry, including probing the impacts of polyploidy and genome size variation on ecological interactions and climate adaptation in this system, as well as elucidating the genetic basis of the physiological tradeoff between water use efficiency and relative growth rate exhibited by the Sonoran desert winter annuals. Making such mechanistic links between genetic changes and ecology is not only a key aim in the field of evolutionary ecology but is a pressing task to evaluate responses to global climate change.

## ACKNOWLEDGEMENTS

The authors thank the Arizona Genomics Institute for their input on the sequencing strategy and methodology. The authors also thank members of the Dlugosch and Barker labs for their feedback on the manuscript. This work was supported by funding from United States National Science Foundation grants #1256792 to D.L.V., #1750280 to K.M.D., and #2022055 support for P.C.N., and from United States Department of Agriculture grants #2023-67013-40169 to K.M.D. and #2024-67012-43394 to J.A.P.

## AUTHOR CONTRIBUTIONS

K.M.D. and L.D.V. conceived of the study. P.C.N. assembled and annotated the reference genome. P.C.N. and J.A.P. conducted the synteny and WGD analyses. P.C.N. wrote the manuscript and J.A.P., K.M.D., and L.D.V. provided feedback. All authors approved the final version of the manuscript.

## DATA AVAILABILITY

The raw genomic sequences (SRR29849658), genome assembly (JBHHJY000000000), and plant tissue sample description (SAMN42375859) are available under the NCBI bioproject PRJNA1133066. Raw transcriptomic sequences (SRR30591463) are linked to the same bioproject. The chloroplast genome is included in the full genome assembly. The structural annotation and functional annotation of the final assembly and chromosomes, as well as relevant data and output from repeat masking, K-mer analysis, OrthoFinder, GENESPACE, and wgd, are archived in a public repository on Zenodo (DOI: 10.5281/zenodo.13851913). Scripts used to generate the assembly and annotation as well as conduct the analyses are available on github (https://github.com/poppynorthing/PERE_genome/).

### Supporting Information

Additional Supporting Information may be found online in the Supporting Information section at the end of the article. Supplementary tables are available in the “Appendix_S1_tables” file and supplementary figures are available in the “Appendix_S2_figures” file.

