## Appendix_S2_figures for "Chromosome-scale reference genome of *Pectocarya recurvata*, a species with one of the smallest genome sizes in Boraginaceae"

Supplemental Figures 1 – 6.

**Supplemental Figure 1A-C.** HiFi Library Quality Metrices for the *P. recurvata* genomic library. A) Distribution of read lengths. B) Distribution of read qualities. C) Number of reads of a given quality and length. M84082_240116_105529_S3 refers to the default library name given by the Revio sequencer (PacBio). HiFi library sequences are available on the sequencing read archive on NCBI (SRR29849658) linked to our NCBI bioproject (PRJNA1133066).

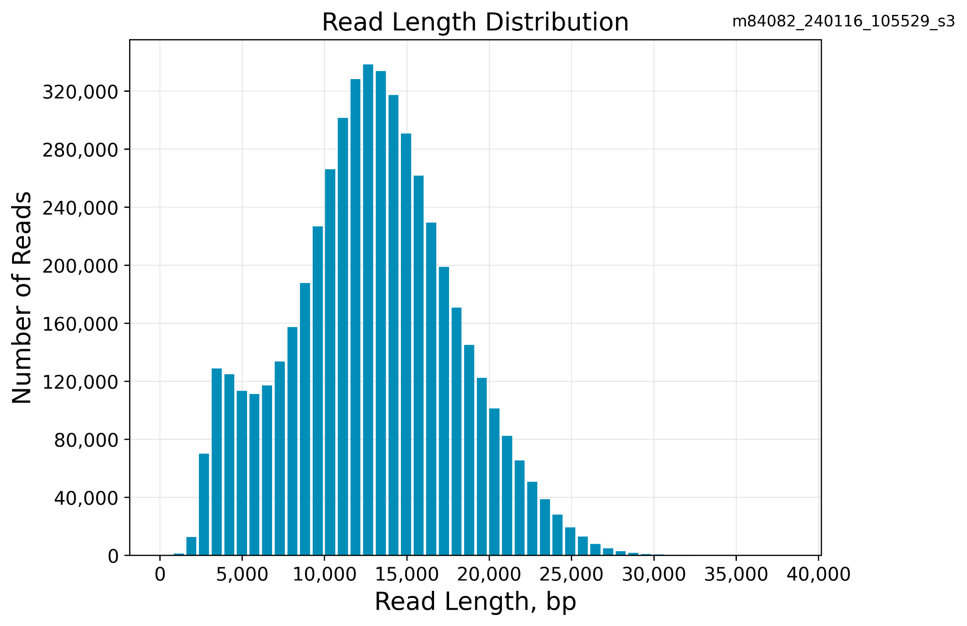

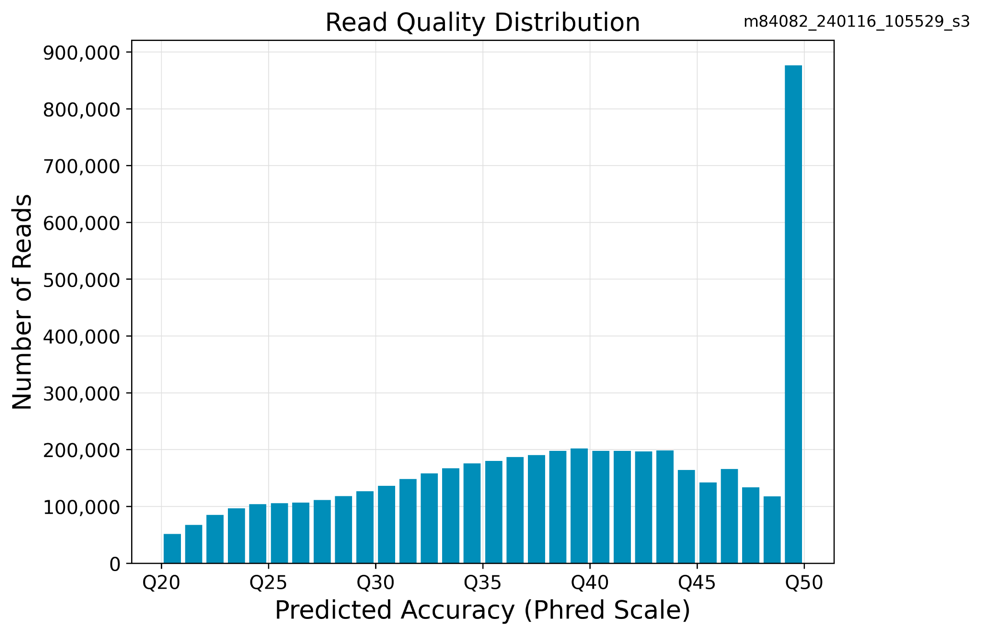

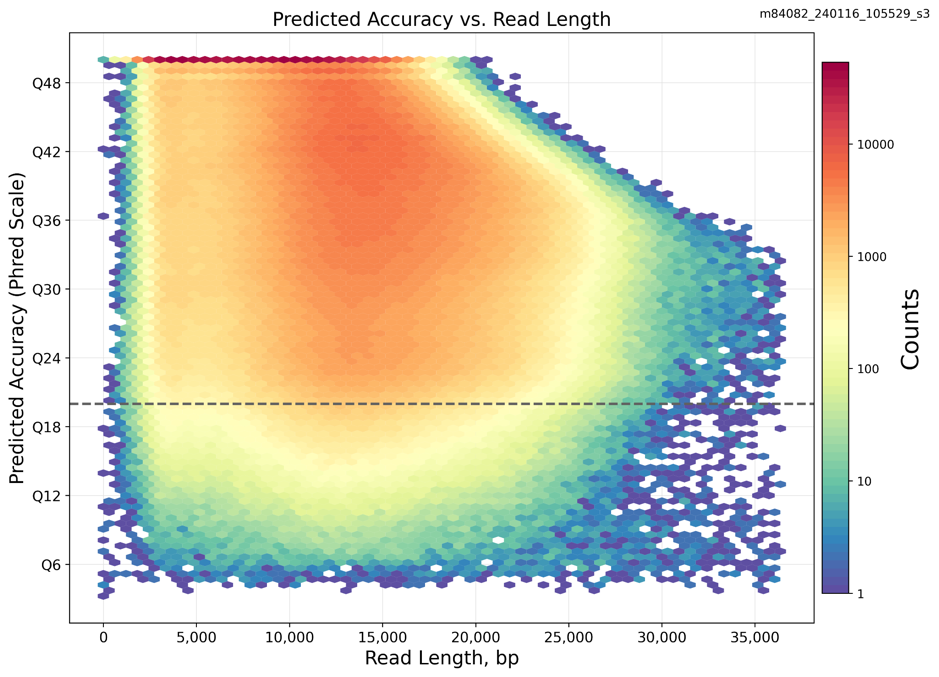

**Supplemental Figure 2A - C**. Genome size and ploidy were estimated by analyzing the distribution and coverage of 17-mers from our HiFi library. A) GenomeScope (Vurture et al., 2017) profile of 17-mers from *P. recurvata*. B) Model output of 17-mers from GenomeScope C) SmudgePlot (Ranallo-Benavidez et al., 2020) of log-transformed coverage of 17-mers indicating that our sequenced individual is a putative tetraploid.

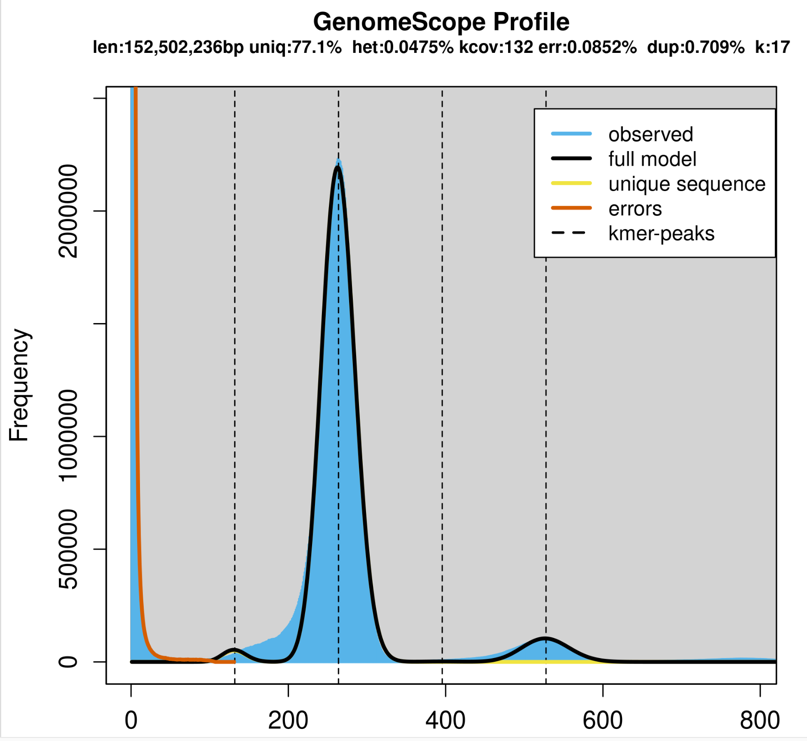

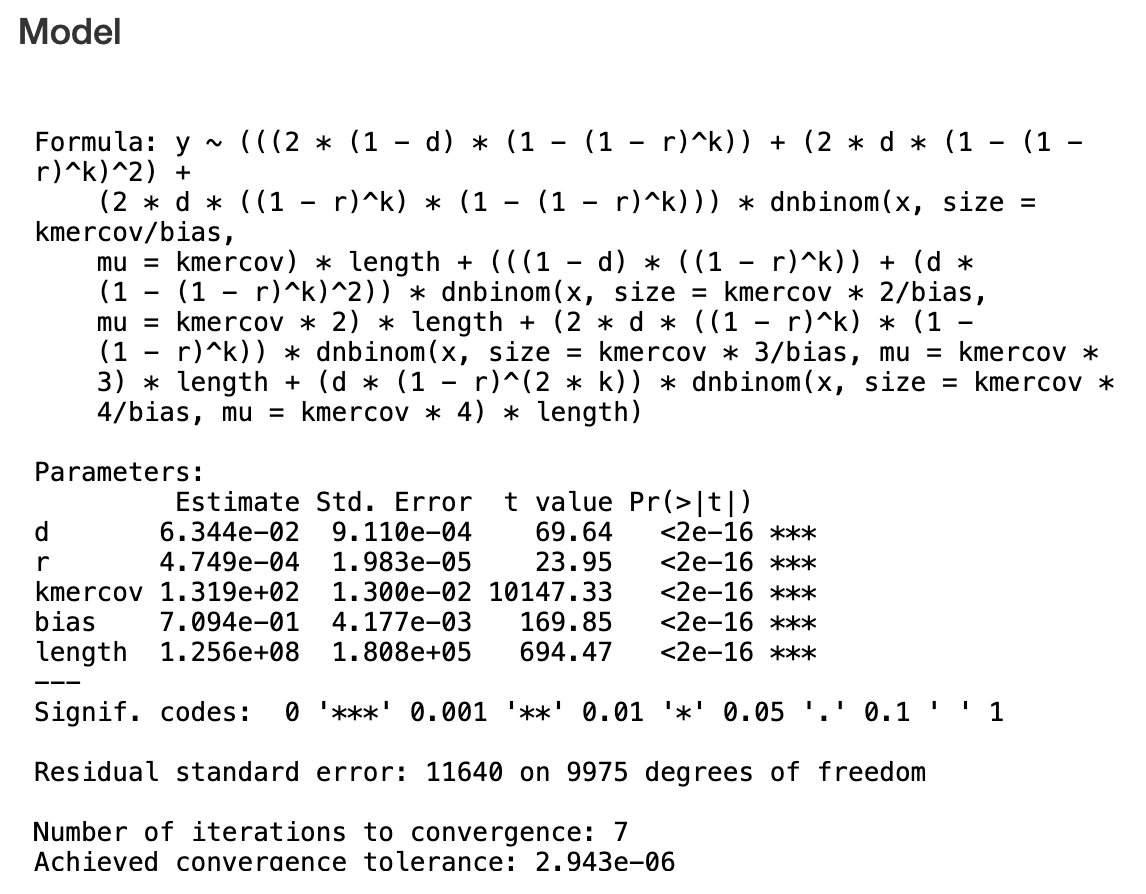

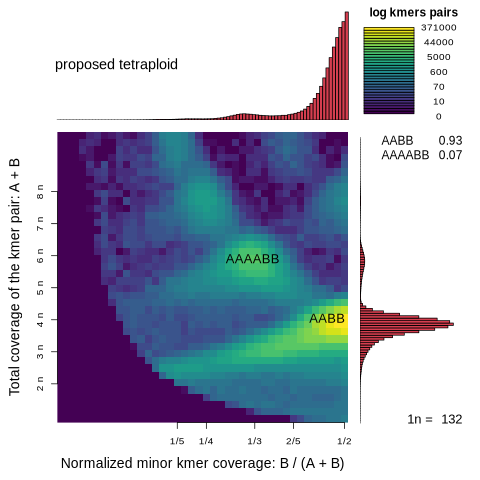

**Supplemental Figure 3A-B.** Self synteny of the *P. recurvata* chromosomes reflects an ancient whole genome duplication that occurred in the evolutionary history of Boraginaceae. A) Self-self dot plot of the raw syntenic hits in the *P. recurvata* chromosomes generated by GENESPACE (Lovell et al., 2022). B) Additional Ks plots to the one included in the paper in Figure 2 (top left panel in this figure) from wgd (Chen et al., 2024). Grey bars reflect the Ks values for *all* paralogs, while the green bars only include Ks values for *syntenic* paralogs.

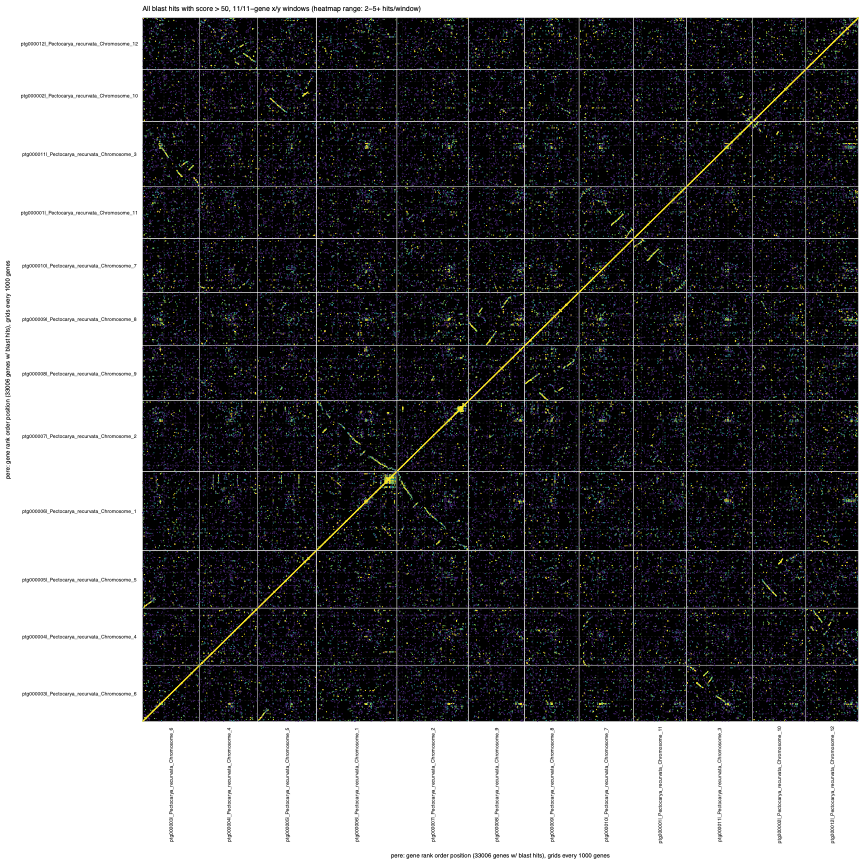

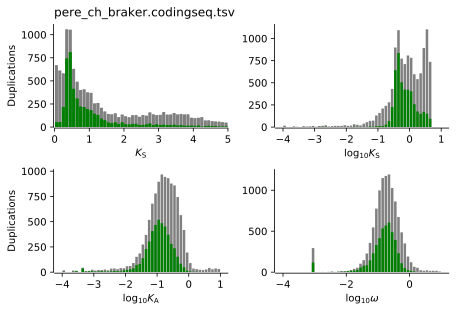

**Supplemental Figure 4A-B.** Self synteny of the *E. plantagineum* chromosomes. A) Self-self dot plot of the raw syntenic hits in the *E. plantagineum* chromosomes generated by GENESPACE (Lovell et al., 2022). B) Ks plots from wgd (Chen et al., 2024); grey bars reflect the Ks values for *all* paralogs, while the green bars only include Ks values for *syntenic* paralogs.

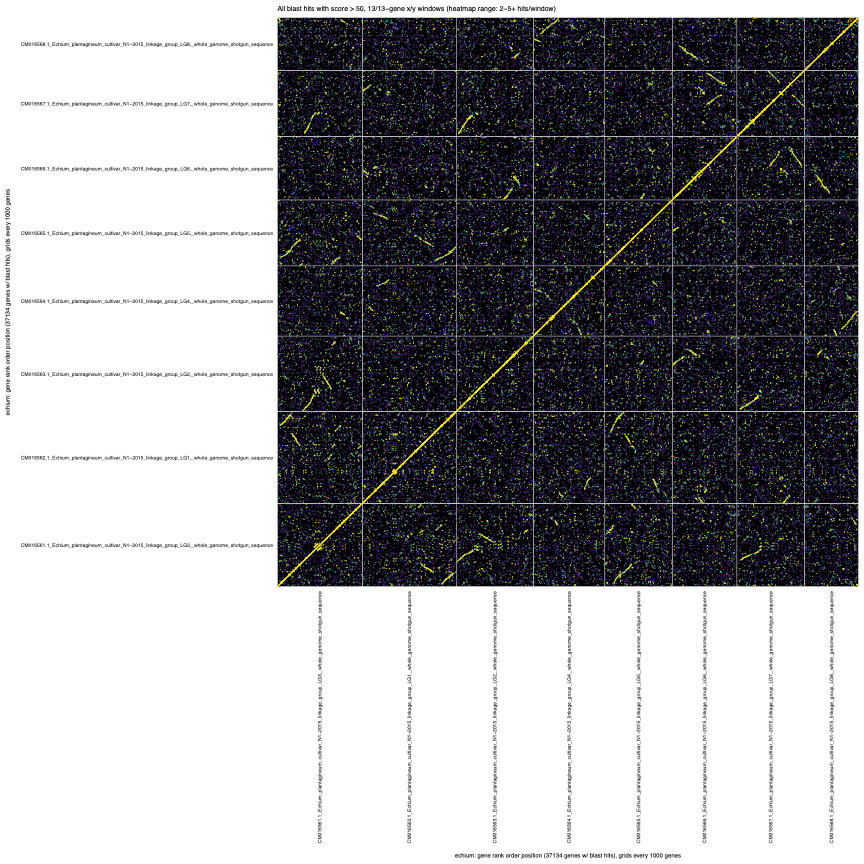

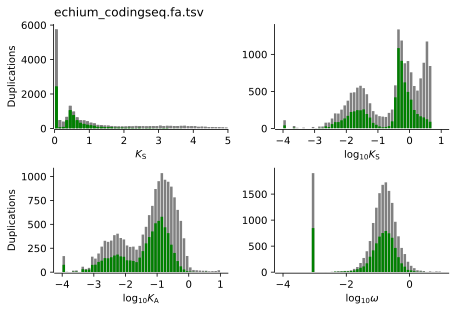

**Supplemental Figure 5.** Dot plot showing only syntenic hits between *P. recurvata* (*n* = 12) and *E. plantagineum* (*n* = 8). *P. recurvata* chromosomes are on the vertical axis, while *E. plantagineum* chromosomes are on the horizonal axis. Color coding indicates syntenic blocks; generated by GENESPACE (Lovell et al., 2022).

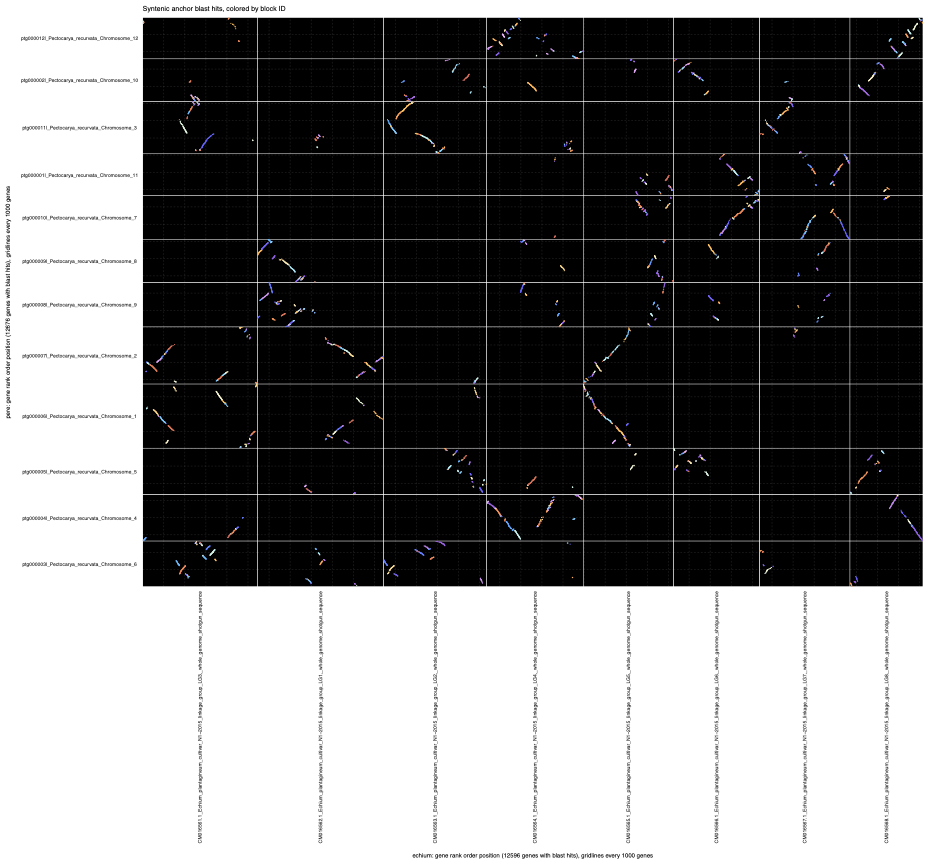

**Supplemental Figure 6.** The syntenic depth ratio of blocks of conserved genes between *P. recurvata* (pere) and *E. plantagineum* (echium) is 2:3. The excess number of duplicate genes in both genomes suggests that these species share a whole genome duplication in their evolutionary history. The excess of triplicate genes in the *E. plantagineum* genome indicates that an additional, independent whole genome duplication has occurred in the history of *E. plantagineum*. Figure generated using pythonic MCscan (Tang et al., 2008b).

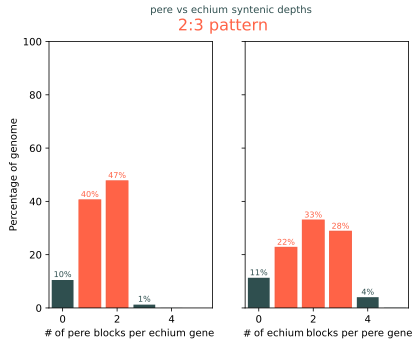

**Supplemental Figure 7.** The annotated *P. recurvata* chloroplast genome of *P. recurvata*.

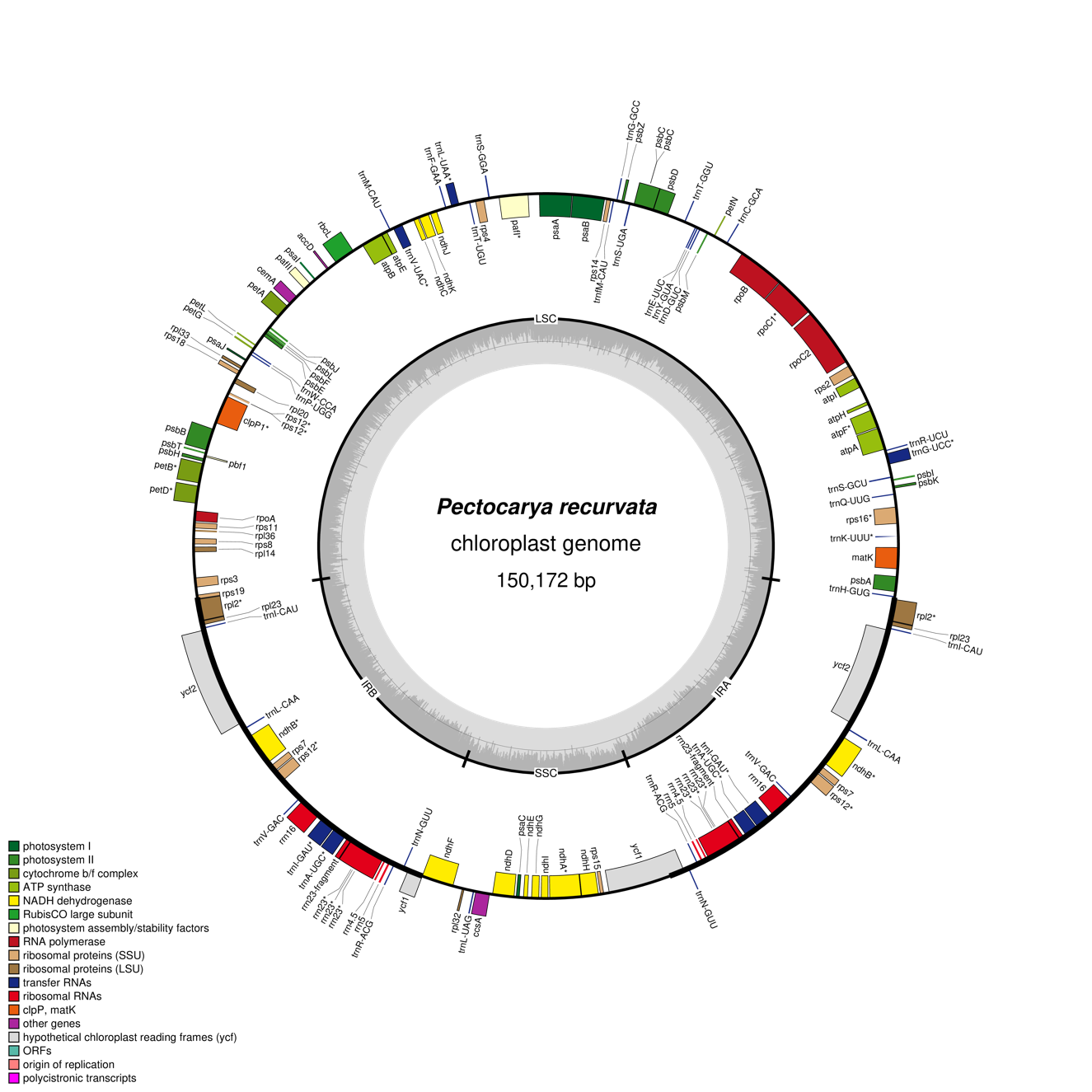
